# CladeOScope-GSA: A Tool for Predicting Functional Gene Interactions via Phylogenetic Profiling

**DOI:** 10.1101/2025.06.03.657590

**Authors:** Maya Braun, Idit Bloch, Dana Sherill-Rofe, Christina Canavati, Elad Sharon, Yuval Tabach

**Author notes:** These authors contributed equally (Maya Braun, Idit Bloch).

## Abstract

Understanding gene function and protein network dynamics is a key challenge in biology and human health. In the light of the exponential growth in genomic data, phylogenetic profiling methods became more powerful in analysing thousands of genomes to uncover evolutionary associations and predict functional interactions [1,2,4,13,14,16,18,23,24,26]. It is based on the well-supported hypothesis that genes that work together are subject to similar evolutionary pressures and therefore their conservation patterns across phylogeny are correlated.

We introduce CladeOScope-GSA, a web-based tool designed for phylogeny-aware analysis of gene sets across extensive comparative genomics datasets. By integrating clade-focused normalized phylogenetic profiling (NPP) with interactive visualization, CladeOScope-GSA enables researchers to identify evolutionary patterns within user-defined gene sets. Building upon the infrastructure of the original CladeOScope visualization platform, the addition of statistical gene set analysis, expanded phylogenetic and genomic data, enhanced visualization, and improved statistical tools represent a significant advancement in the tool’s capabilities. We demonstrate its utility through case studies spanning 1905 species, showcasing how CladeOScope-GSA empowers researchers to extract biological insights from complex comparative datasets.

The web tool is accessible at https://tabachlab.shinyapps.io/CladeOScope

## Introduction

Phylogenetic profiling (PP) is a powerful method for inferring functions of uncharacterized genes through comparative genomics [9,14,22]. A gene’s phylogenetic profile represents its conservation and loss patterns across a set of genomes [14]. PP is founded on the robust hypothesis that genes sharing similar phylogenetic profiles are likely to interact functionally [8,14,23,24].

Initially, PP employed binary scores to indicate the presence or absence of a gene across species [7,10,14]. Over time, more sophisticated approaches like normalized phylogenetic profiling (NPP) were developed to account for varying levels of conservation, as proteins are often not fully lost but exhibit significant divergence [19,23–25]. NPP employs a continuous conservation metric normalized by the phylogenetic distance from the query species (in our case, human).

In recent years, we found that a signal of co-evolution within clades (groups of species sharing a common ancestor or segments of the evolutionary tree) might indicate functionally related genes. This “clade-wise” NPP approach has been successfully applied to uncover novel DNA repair genes [18,21] and identify potential therapeutic targets for MECP2 [26] and ACE2-associated disorders [4]. Moreover, a “guilt by association” approach enabled the discovery of disease-causing genes implicated in rare genetic diseases [5].

CladeOScope [25] was initially designed to enable users to investigate single genes, identifying co-evolved genes across all eukaryotes and within specific clades. By leveraging clade-based phylogenetic analysis, it provided a user-friendly, interactive webtool for this purpose. Few tools currently offer comparable functionality [6,12,16]. CladeOScope-GSA introduces the capability to analyze gene sets, presenting the conservation patterns of multiple input genes/proteins across 1905 genomes. It uncovers significant co-evolution patterns at various evolutionary scales, providing insights into the biological contexts of novel genes and facilitating the identification of co-evolution trends within predefined gene groups.

### Multiple gene (gene set) analysis

Multiple gene analysis is designed to evaluate phylogenetic correlations among genes in a user-defined set. Initially, CladeOScope-GSA assesses the co-evolutionary significance of the gene set using two distinct metrics (see Materials and Methods). Upon submission, the query set is compared within each clade against 1000 pre-calculated random gene sets of the same size. Additionally, the query is compared to a positive control set (Krebs cycle genes according to KEGG, Tabach 2013 [24]). If the query set exceeds the size of the Krebs cycle gene set, random genes are added to the control set to match its size. This analysis provides users with insights into the degree of co-evolution within their gene set. CladeOScope-GSA generates NPP and correlation heatmaps alike to single gene analysis. Notably, non-conserved genes are excluded from both the correlation heatmap and the random significance analysis.

### Clade-wise analysis of porphyria genes reveals evolutionary patterns linked to disease etiology

Porphyria comprises metabolic disorders caused by mutations in heme biosynthesis genes, leading to enzyme deficiencies and toxic porphyrin accumulation [27]. Analysis of the nine porphyria-related genes reveals distinct conservation patterns (Fig. 1). Nematodes and certain fungi, which are unable to synthesize heme de novo and must obtain it externally [15], exhibit low to absent conservation of these genes, leading to their distinct clustering. This co-evolutionary pattern is significant across eukaryotes, nematodes, and fungi, but not in chordates or other clades.

**Figure 1:**
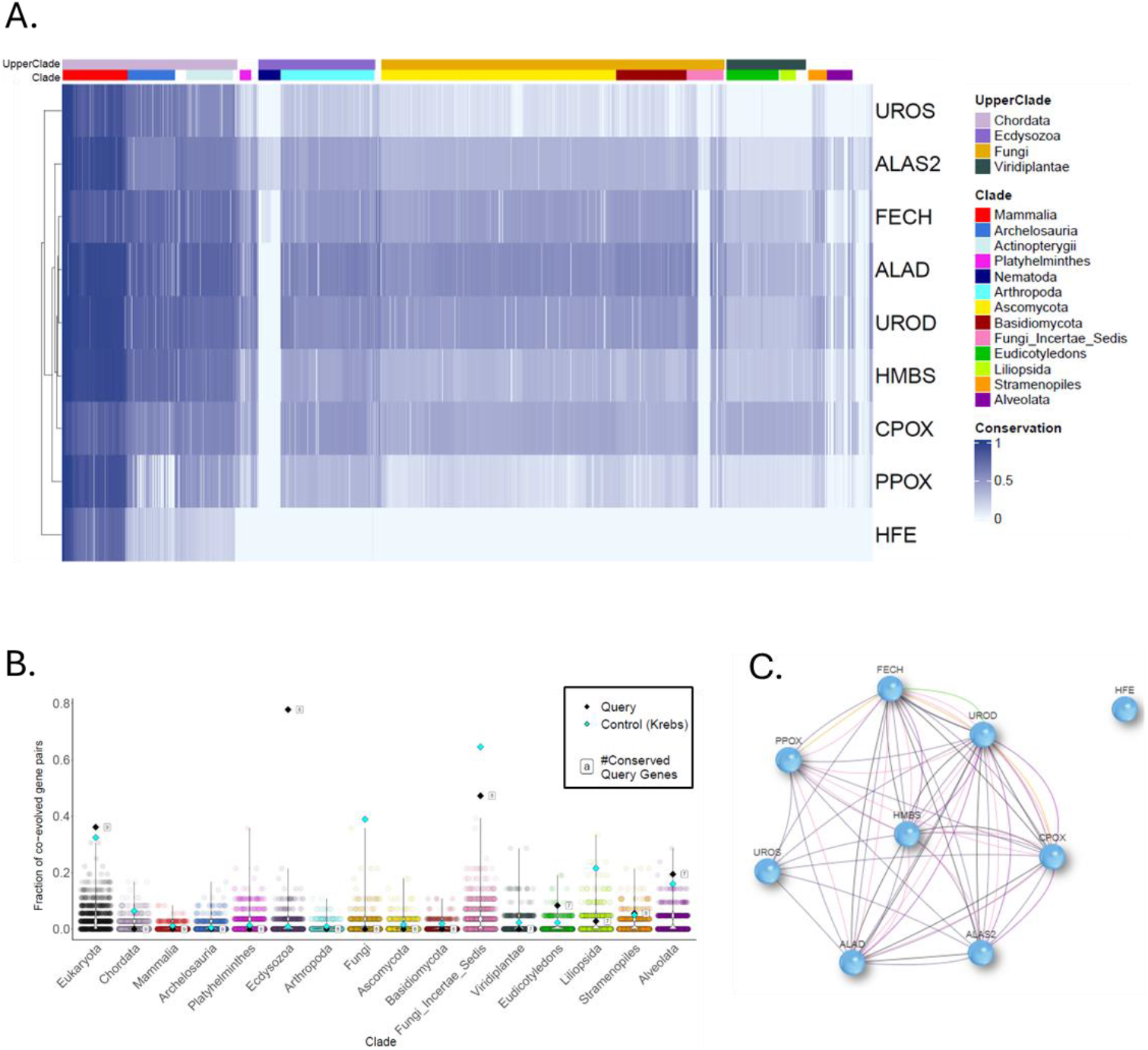
Porphyria genes exhibit strong co-evolution, evident in the similarity of their phylogenetic profiles. (A) Each row corresponds to a gene, and each column represents one of 1,905 species. Dark blue indicates high conservation of the gene within the species, while white signifies the absence of an ortholog for the human query gene in that species. (B) demonstrates how significantly co-evolved the query gene set is in eukaryotes, ecdysozoa, and fungi, in comparison to thousands of random sets of the same size, and to the Krebs cycle gene set. Each dot represents a set of random genes, colored by the clade in which the set was examined. Lastly, (C) the co-evolution network illustrates the genes that co-evolve with one another.

Interestingly, HFE (Homeostatic Iron Regulator), a relatively newly evolved protein, does not cluster with the porphyria-related genes. While most subtypes of porphyria arise from direct enzyme defects, a specific subtype is indirectly linked to iron overload caused by HFE gene mutations. HFE variants, associated with hereditary hemochromatosis, exacerbate iron overload, inhibiting uroporphyrinogen decarboxylase and thus triggering porphyria cutanea tarda [11]. These findings underscore the functional connections among porphyria-related genes and highlight the potential of clade-wise NPP analyses to uncover disease mechanisms and etiology.

### Phylogenetic profiling of dynein gene families reveals structural subgroup clustering

The second example focuses on the dynein gene group (Fig. 2). These genes were analyzed based on the dynein gene set as appears in the HGNC database [17]. The phylogenetic profile of dynein genes reveals distinct clustering based on their functional and structural subgroups. The heatmap highlights patterns of conservation across clades, with specific clustering observed among eukaryotes, ecdysozoa, fungi, and plants. Functional subgroups display co-evolutionary trends, suggesting shared evolutionary pressures and structural roles among dynein gene families [3].

**Figure 2:**
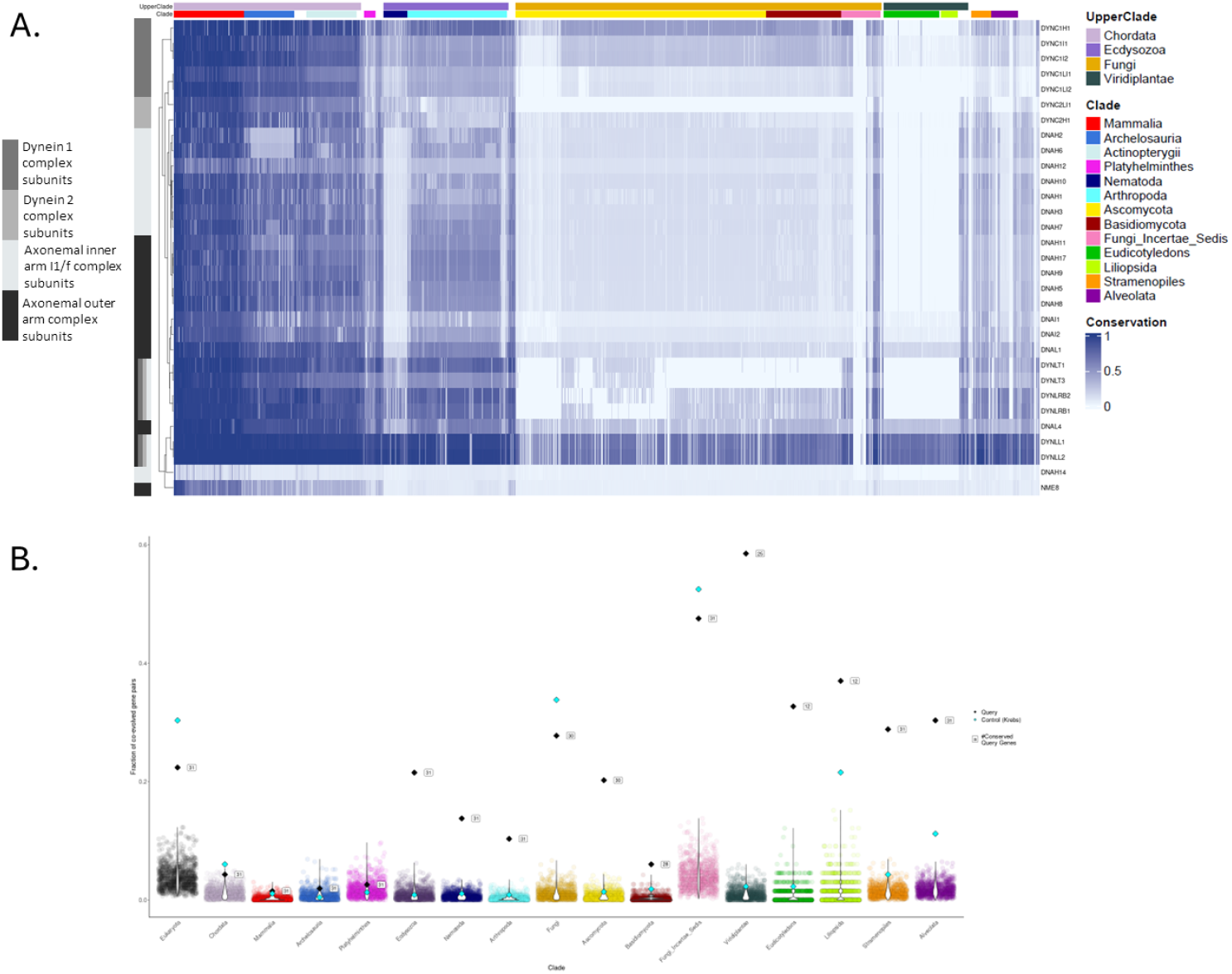
The pylogenetic profile of dynein genes clusters by their structural subgroups. (A) Each row represents a gene, and each column corresponds to one of 1,905 species. Dark blue indicates strong conservation of the gene in a given species, while white denotes absence of an ortholog to the human reference gene. The bar to the left of the heatmap indicates the structural subgroup classification of each dynein gene. (B) illustrates the extent to which the query gene set is significantly co-evolved in eukaryotes, ecdysozoa, fungi, and plants, compared to thousands of randomly generated gene sets of the same size and to the Krebs cycle gene set. Each dot represents a random gene set, colored according to the clade in which it was analyzed.

### Summary and Perspectives

CladeOScope-GSA introduces a significant extension to phylogenetic profiling by enabling clade-aware analysis of gene sets. This approach captures co-evolutionary signals and lineage-specific conservation patterns. Its statistical framework supports rigorous assessment of gene set enrichment or depletion across diverse taxa, facilitating insights into functional relationships, evolutionary constraints, and disease mechanisms.

By combining analytical power with a freely available user-friendly, web-based interface, CladeOScope-GSA makes high-resolution phylogenomic analysis broadly accessible. Its application across various case studies highlights its utility as a versatile tool for advancing research in comparative genomics, functional annotation, and evolutionary biology.

## Materials and Methods

### Updated database content and statistics

The normalized phylogenetic profiling (NPP) matrix was updated to include 19,888 human genes across 1,905 eukaryotic species genomes, following a methodology similar to that previously described [2,4,16,23–25], with the following modifications:

#### 1. Genome Databases

The query Homo sapiens proteome was downloaded from UniProt proteomes on 17.3.2020. Reference whole proteomes were retrieved from three databases: Ensembl release 100, NCBI Genomes Refseq (August 2020), and Uniprot “reference proteomes” Release 2020_04. Proteome sequences for each species were merged, and duplicate sequences were filtered, resulting in a dataset of 1,905 species.

#### 2. Gene Filtering

To normalize the matrix, each gene’s ortholog bitscore was divided by the bitscore of its human self-hit to account for protein length, creating a Length Normalized Phylogenetic Profiling (LNPP) matrix. Genes with zero values across all species (43 in total) were excluded, resulting in a matrix of 20,355 genes and 1,905 species.

The number of genes was further reduced to 19,888 to align with the GeneCards gene set. Gene symbols were standardized according to GeneCards nomenclature to enable seamless integration with its platform.

### Evaluating Co-Evolution Significance: Threshold Score or Cluster Score

Threshold Score - The threshold score measures the percentage of pairwise Pearson correlations within a query gene set that exceed predefined thresholds (0.65–0.9). For a gene set of size N, the total number of gene-gene correlations is (n^2^/2)-n. CladeOScope computes the Pearson correlation (ranging from -1 to 1) and calculates the percentage surpassing the chosen threshold. While paralogous genes often exhibit correlations ≥0.9, and random genes around 0, non-paralogous genes that are known to be co-evolved may show values around 0.7. Therefore, a threshold value around 0.7 is recommended.

Cluster Score - In some cases, a gene set comprises several gene clusters that do not correlate with one another. In such cases, the correlation percentage exceeding some threshold may be low (because many genes do not correlate with each other), even though there are sub-groups that are co-evolved. The cluster score addresses this problem and therefore tends to be more relevant in larger (>50) gene sets, where not every gene is co-evolved with another gene. To compute this score, CladeOScope performs hierarchical clustering (complete linkage) of the gene set. The dendrogram is then “cut” at a height of 0.2 (indicating a minimal correlation of 0.8). At this height, CladeOScope calculates the number of clusters, their sizes, and uses these measures to compute the score:

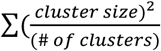

where “cluster size” is the number of genes clustered together in each cluster and “# of clusters” (number of clusters) is the number of independent clusters at a height of 0.2.

### Enhanced web interface

The website interface was upgraded to enhance usability and data visualization. Key improvements include detailed clade annotations displayed atop heatmap columns, improved color schemes and proportions in heatmaps and tables, and an interactive network representation of phylogenetic profile correlations, displaying interactions exceeding a threshold of 0.7.

### Collaboration with the Genecards suit

We recently launched a collaboration with GeneCards [20] enabling direct integration between the two platforms. Each gene in CladeOScope now links to its corresponding GeneCards page. Additionally, the GeneCards page for each gene lists its top 100 co-evolved genes across 17 clades and all eukaryotes, with direct links to relevant heatmaps on the CladeOScope website.

